## Supplementary Figures for "Deep convolutional and conditional neural networks for large-scale genomic data generation"

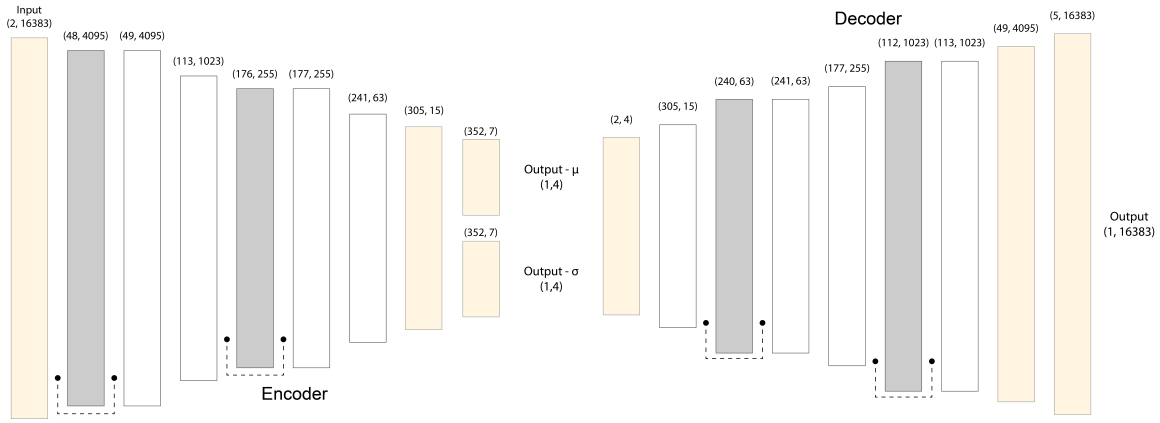


**Figure S1. Architecture of the variational autoencoder (VAE) model for the 10,000 SNP dataset.** Generic blocks of the encoder and decoder (white rectangles) are conceptually the same with the generic critic and generator blocks respectively (Fig 4a), except that there are no latent space channels concatenated to the input and no additional noise vectors at each block. The major difference from WGAN in terms of architecture is the last block of the encoder, which encodes mu and sigma as the mean and the standard deviation of the generated distribution, which are used to sample the latent space. Dotted connections are residual connections where the input value is added to the output value of the block before passing to the next block. Numbers in parentheses above blocks show channels and length, respectively (C, L).

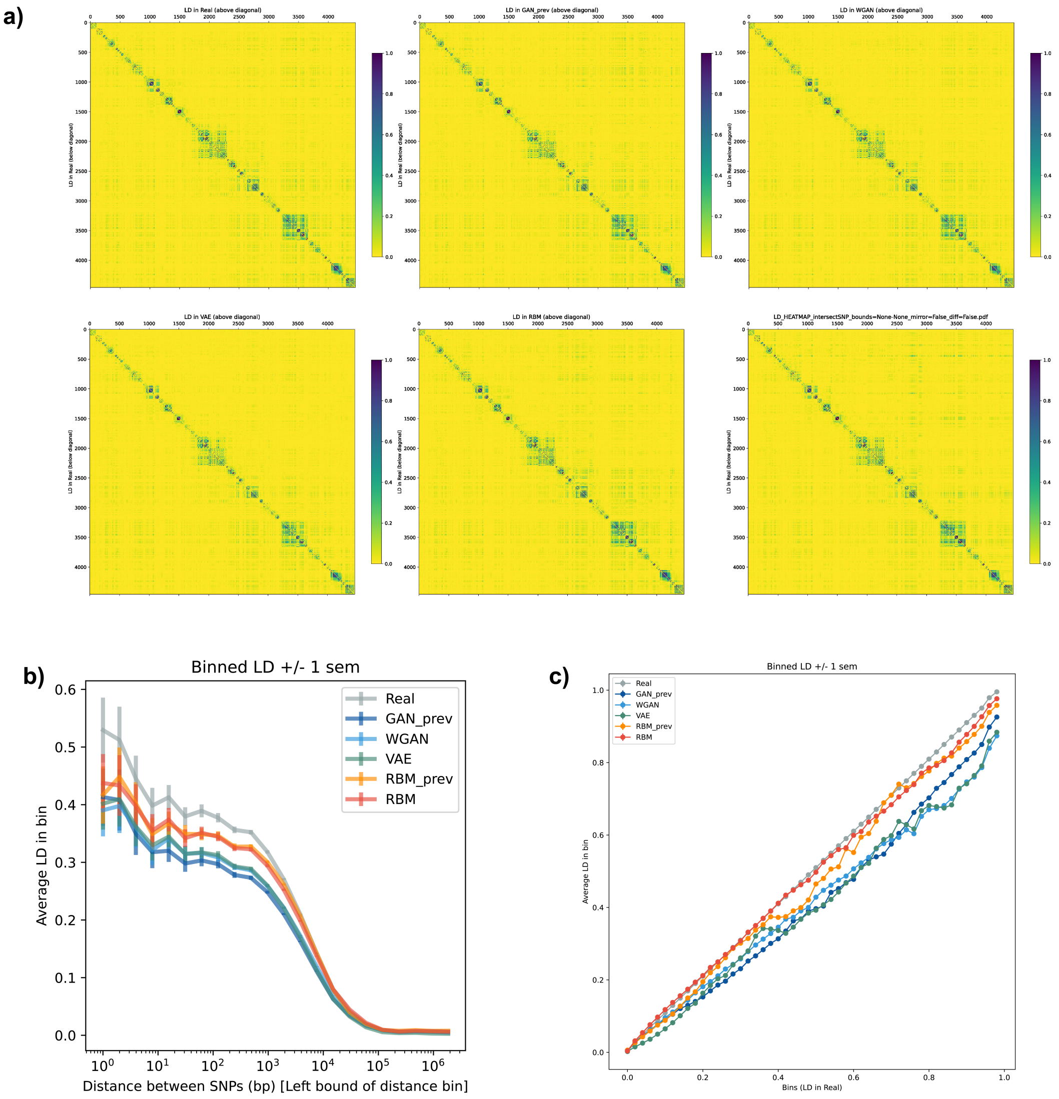


**Figure S2. Comparative linkage disequilibrium (LD) analysis of artificial genomes generated by different models for the 10,000-SNP data. a)** LD heatmap based on $r^2$ matrices. Sections below diagonals correspond to LD in real genomes and sections above diagonals correspond to LD in artificial genomes. **b)** LD decay as a function of SNP distance. SNPs were binned based on distance and average LD was calculated. **c)** LD decay correlation between real and artificial datasets. x axis corresponds to real LD bins and y axis corresponds to generated LD bins. Sites fixed in any of the datasets were removed for all the LD calculations.

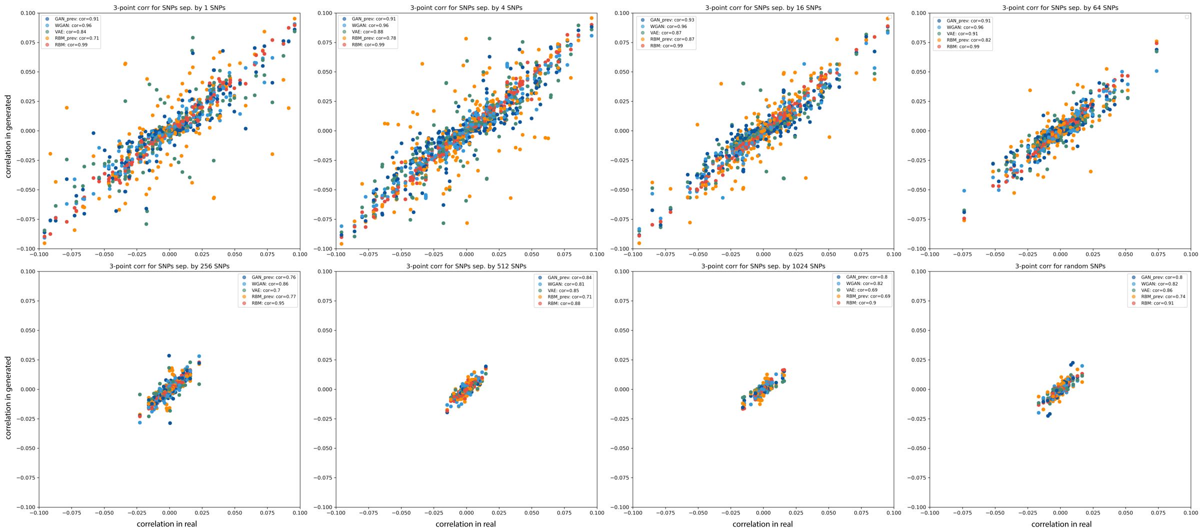


**Figure S3. 3-point correlation analysis of SNP triplets for the 10,000-SNP data with inter-SNP distances of 1, 4, 16, 64, 256, 512 and 1024 (from left to right, top to bottom).** The last panel (bottom right) shows correlation for triplets of SNPs drawn randomly. In each plot, drawing order (z-order) of each AG group is shuffled.

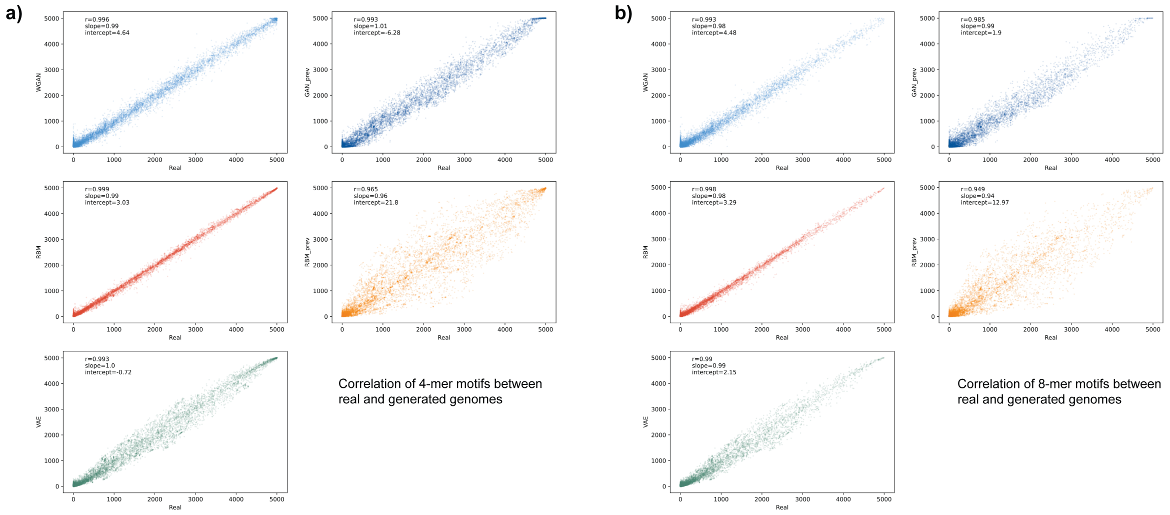


**Figure S4. Analysis of small haplotype motifs between real and generated 10,000-SNP datasets** for **a)** 4-mer and **b)** 8-mer non-overlapping windows. For each unique k-mer in each window, number of occurrences in the real dataset was compared to the same number in the AG dataset. Each point corresponds to the occurrence number in real (x-axis) and AG (y-axis) datasets. Values presented inside the figures are Pearson's r, ordinary least squares regression slope and intercept.

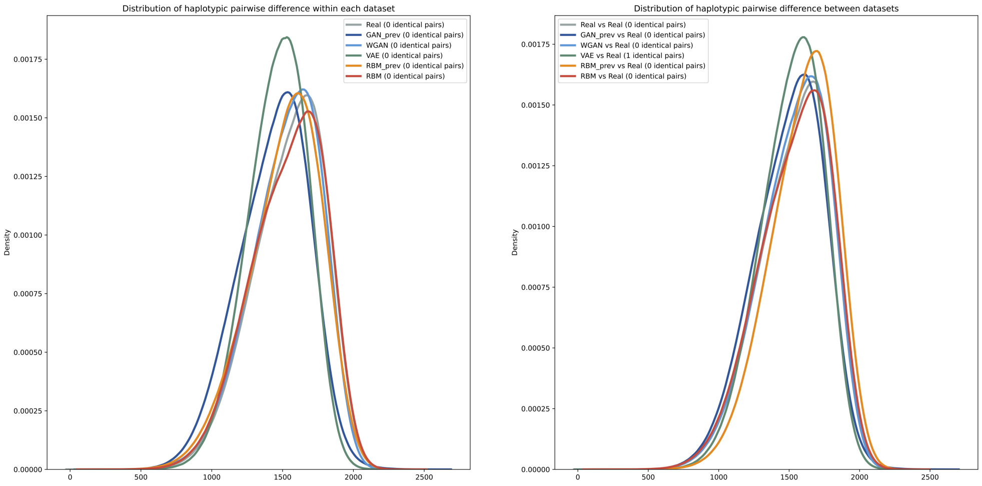


**Figure S5. Distribution of haplotypic pairwise difference within (left figure) and between (right figure) 10,000-SNP datasets.**

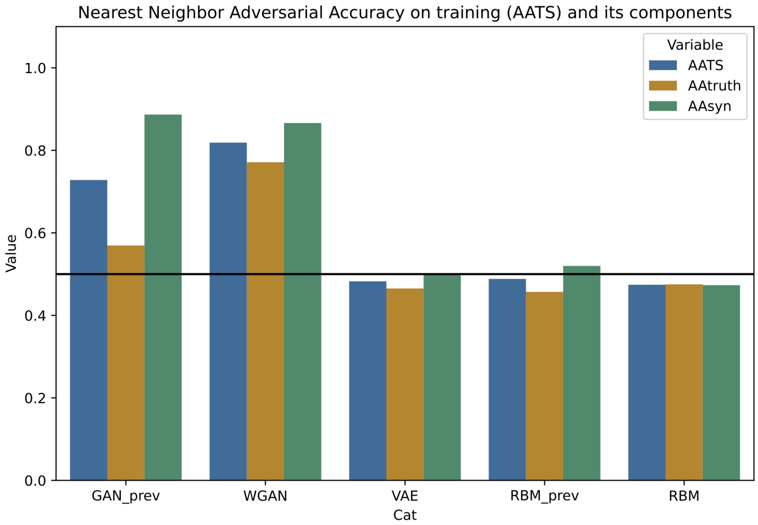


**Figure S6. Nearest neighbour adversarial accuracy (AATS) of artificial genomes generated by different models for the 10,000-SNP dataset.** Values below 0.5 (black line) indicate overfitting and values above indicate underfitting. See Materials and Methods for the details of the metrics.

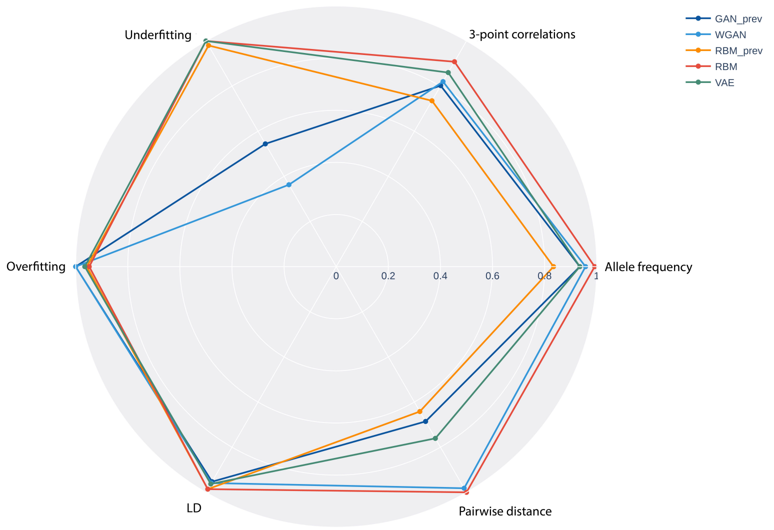


**Figure S7. Radar plot comparing artificial genomes generated by different models for the 10,000-SNP dataset.** Values closer to 0 indicate poor performance whereas values closer to 1 indicate good performance. See Materials and Methods for the details of the representative statistics.

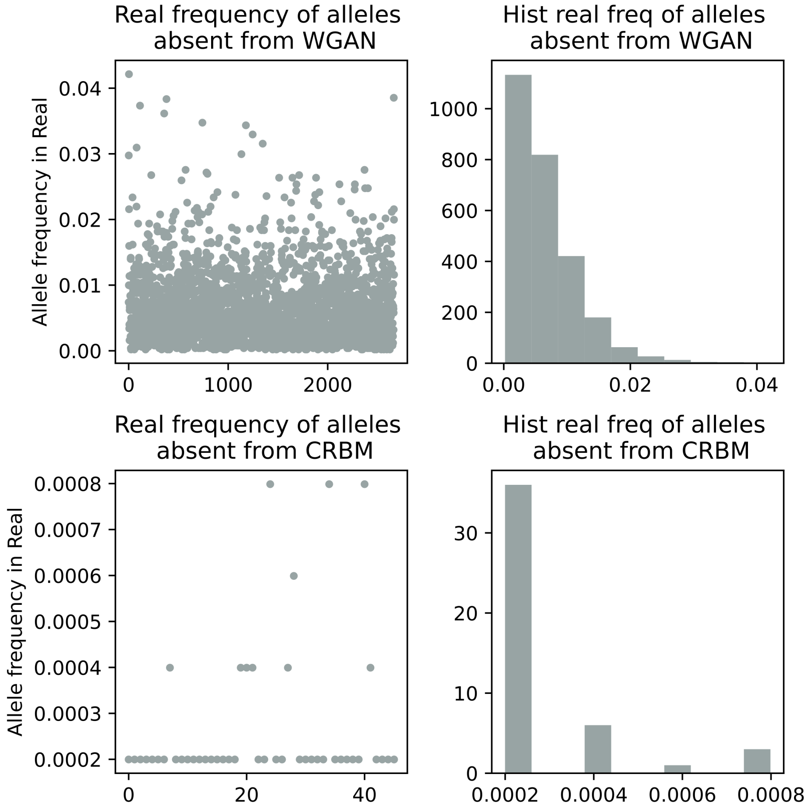


**Figure S8. Analysis of fixed alleles in artificial genomes with 65,535 SNPs.** Left figures show the number of fixed alleles in artificial genomes (x axis) versus the frequency of these alleles in the real dataset (y axis). Right figures show the distribution of the frequency of alleles fixed in the artificial dataset but not fixed in the real dataset.

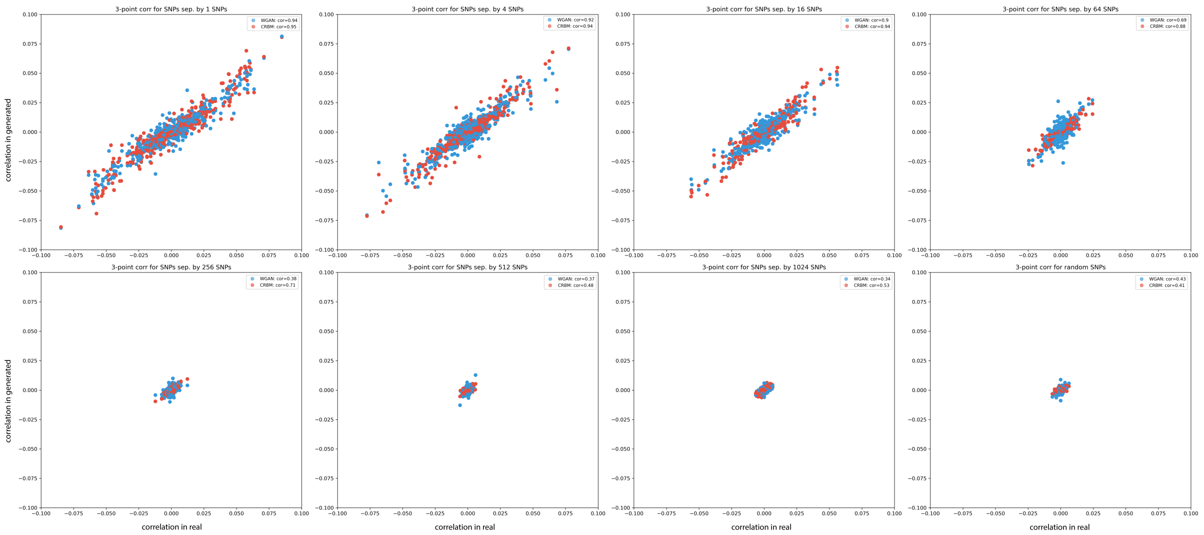


**Figure S9. 3-point correlation analysis of SNP triplets for the 65,535-SNP data with inter-SNP distances of 1, 4, 16, 64, 256, 512 and 1024 (from left to right, top to bottom)** for WGAN and CRBM AGs. The last panels (bottom right) shows correlation for triplets of SNPs drawn randomly. In each plot, drawing order (z-order) of each AG group is shuffled.

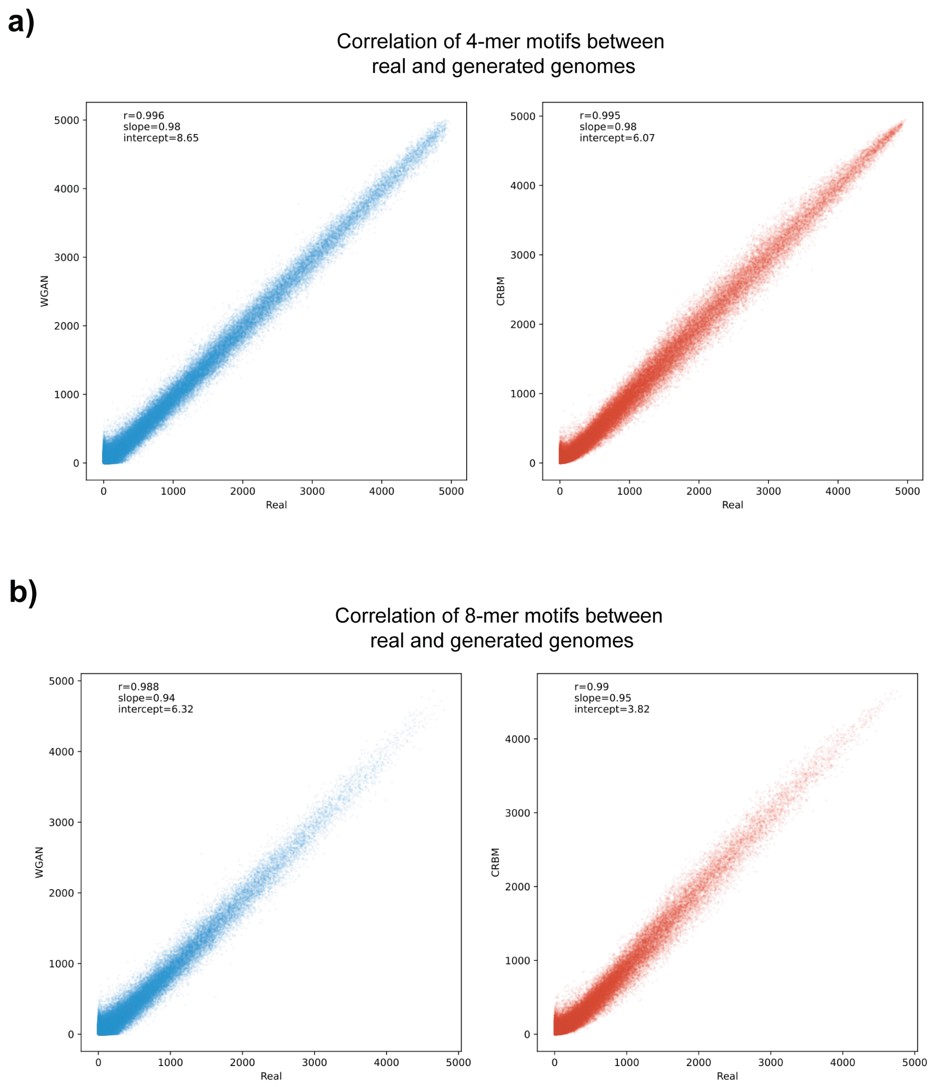


**Figure S10. Analysis of small haplotype motifs between real and generated 65,535-SNP datasets** for **a)** 4-mer and **b)** 8-mer non-overlapping windows. For each unique k-mer in each window, number of occurrences in the real dataset was compared to the same number in the AG dataset. Each point corresponds to the occurrence number in real (x-axis) and AG (y-axis) datasets. Values presented inside the figures are Pearson's r, ordinary least squares regression slope and intercept.

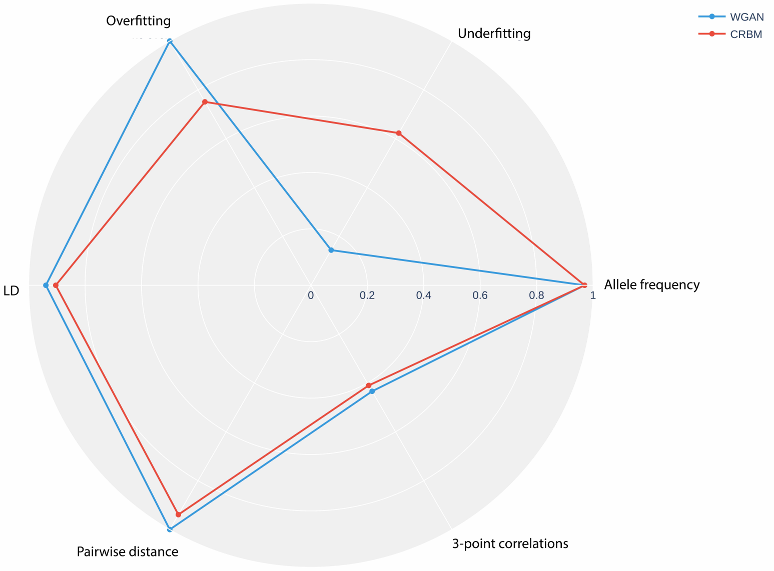


**Figure S11. Radar plot comparing artificial genomes generated by WGAN and CRBM models for the 65,535-SNP dataset.** Values closer to 0 indicate poor performance whereas values closer to 1 indicate good performance. See Materials and Methods for the details of the representative statistics.

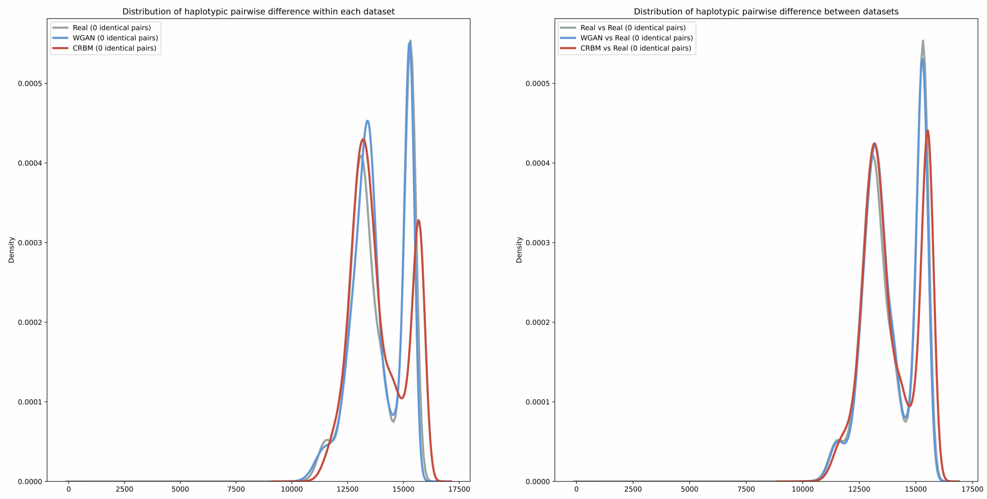


**Figure S12. Distribution of haplotypic pairwise difference within (left figure) and between (right figure) 65,535-SNP datasets.**

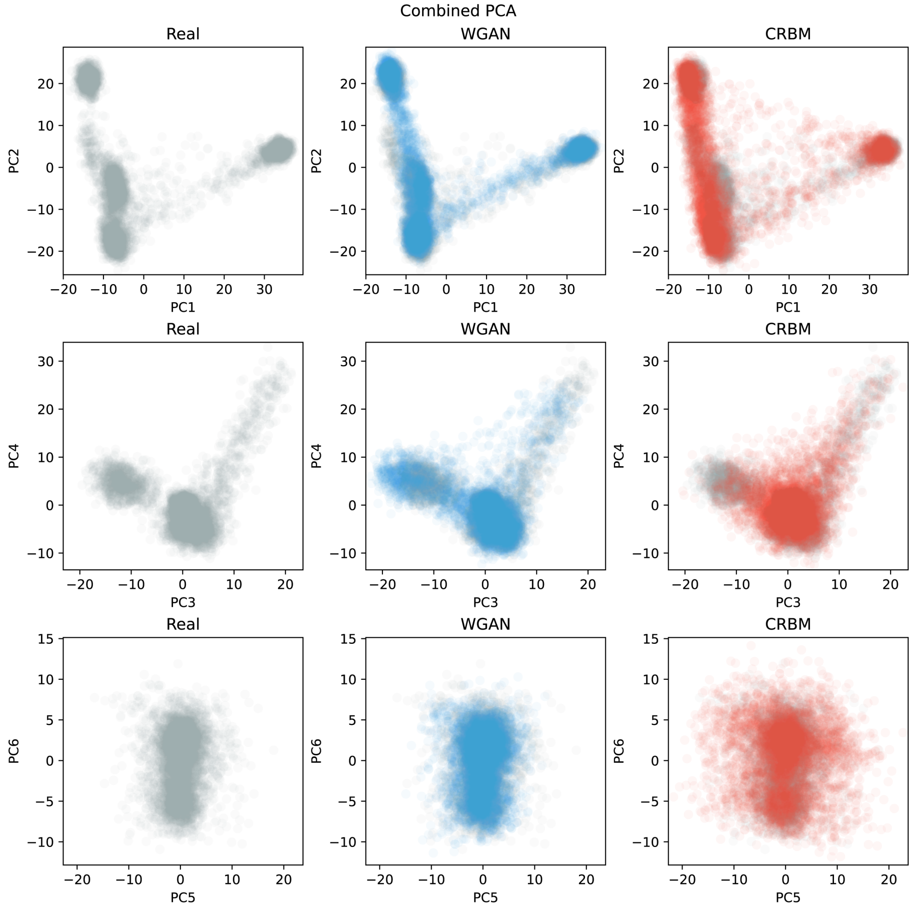


**Figure S13. Principal component analysis (PCA) of combined real and artificial genomes with 65,535 SNPs.**

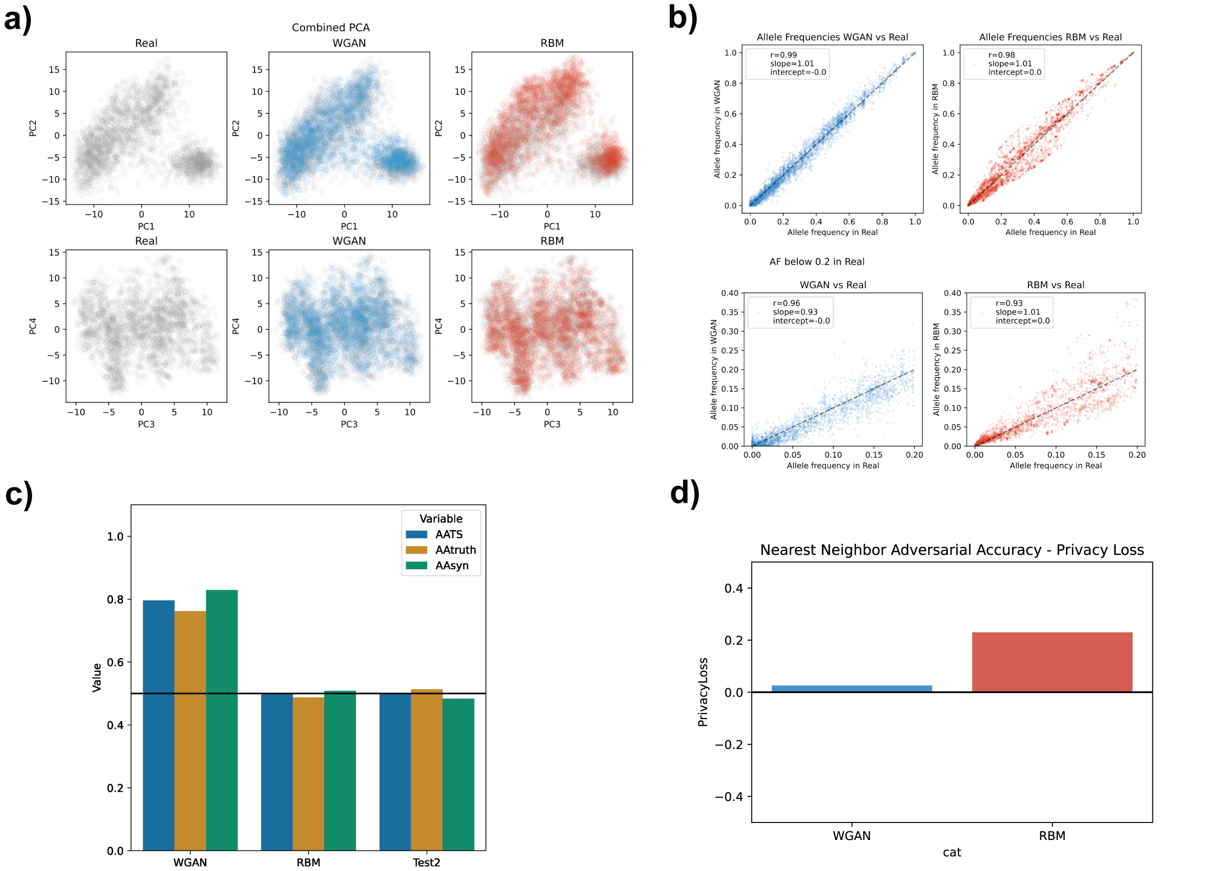


**Figure S14. Analysis of WGAN and RBM generated AGs using 2504 samples from 10,000-SNP dataset. a)** Principal component analysis (PCA) of combined real and artificial genomes. **b)** Allele frequency correlation between real (x-axis) and artificial (y-axis) genome datasets. Bottom figures are zoomed at low frequency alleles (from 0 to 0.2 overall frequency in the real dataset). Values presented inside the figures are Pearson's r, ordinary least squares regression slope and intercept. **c)** Nearest neighbour adversarial accuracy (AATS) of artificial genomes generated by different models and the test set. Values below 0.5 (black line) indicate overfitting and values above indicate underfitting. **d)** Privacy score for WGAN and RBM generated AGs. Values close to 0 indicate no privacy leakage.

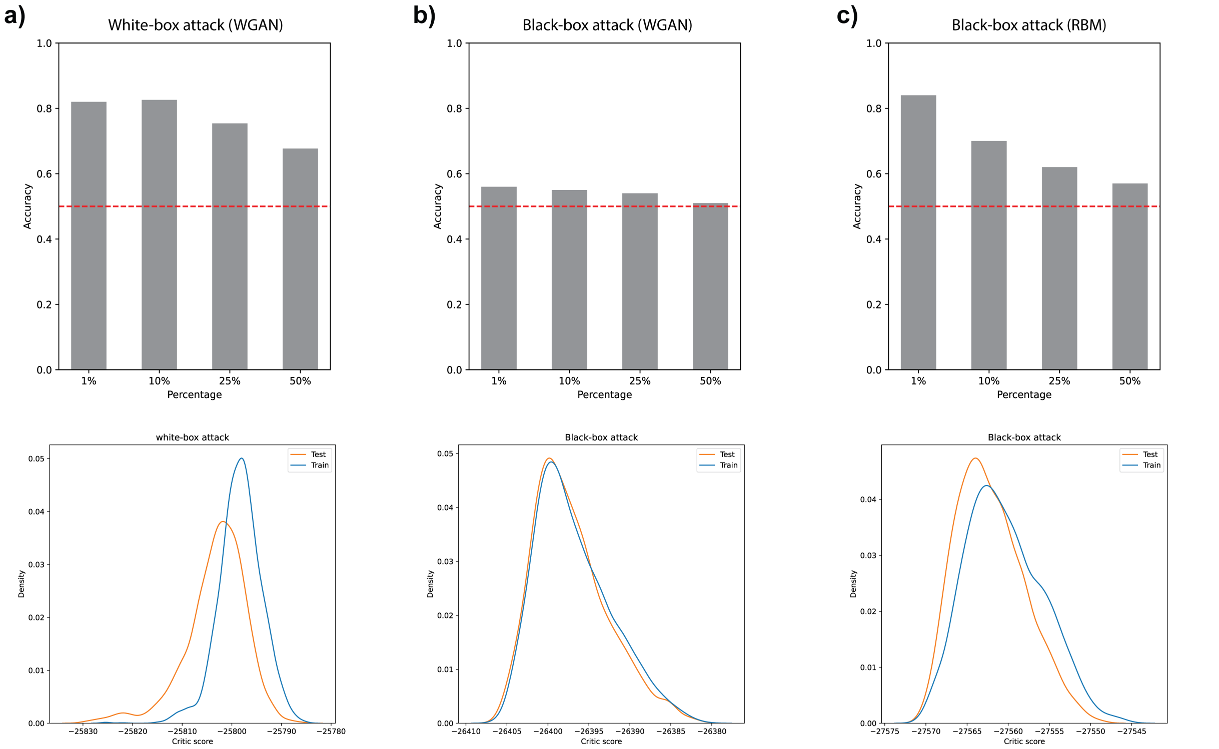


**Figure S15. Membership inference attack on generated 10,000-SNP genomes. a)** White-box attack (adversary has access to the model architecture and weights) on WGAN AGs and black-box attacks with auxiliary information (adversary has only access to the model architecture) on **b)** WGAN and **c)** RBM AGs. For all attacks, the adversary is assumed to know the size of the training set (2504 in this analysis) and possesses a set of samples (5008 in this analysis) suspected of belonging to the training data. For each attack, the critic scores the samples and the adversary sets a threshold for assigning the top $n$ scoring samples to the training dataset. Figures in the upper row show the accuracy of attacks depending on these thresholds (assigned samples ranging from the top 1% to the top 50%). The red dashed lines indicate the accuracy if the $n$ samples were chosen randomly and not based on their scores. Figures in the lower row show the distribution of the critic score for train and test datasets. See Materials and Methods for more details.
